# Compositional selection of phospholipid compartments in icy environments drives the inheritance of encapsulated genetic information

**DOI:** 10.1101/2025.04.16.648126

**Authors:** Tatsuya Shinoda, Natsumi Noda, Takayoshi Watanabe, Kazumu Kaneko, Yasuhito Sekine, Tomoaki Matsuura

## Abstract

The lipid world hypothesis proposes that both intracellular components *and* the chemical composition of the membrane compartment carry heritable information that contributes to protocellular fitness. However, there are few experimental demonstrations of membrane compositional selection and no study has shown that phospholipids—the primary membrane components of modern cells—can support selection of this type. As lipid compartments generally grow in size before fission, faster-growing compartments should enjoy a selective advantage that favors both the membrane itself and the contents it hosts. When situated in icy environments undergoing cycles of freezing and thawing (F/T), we found that the growth of phospholipid vesicles depends on their phospholipid composition: Vesicles with more unsaturated bonds in the acyl chain showed higher growth efficiency. When vesicles composed of lipids with either one or two unsaturated bonds were mixed and subjected to F/T cycles, a selective enrichment of the lipid with two unsaturated bonds was observed. Moreover, selection acting on lipid composition was propagated to the encapsulated genetic material, which was also enriched while selectively neutral. We conclude that phospholipid composition can be a direct target of selection for grown vesicles under an icy environment leading to indirect but concurrent enrichment of compartmentalized genetic molecules.

## Introduction

Compartmentalization was likely a key factor for the origin of life. With distinct compartments that shield internal chemical components from the external environment^1,2^, primordial cells could diversify and compete, leading to early Darwinian evolution^3^. Experimental studies support this view, demonstrating that diversity among encapsulated systems can induce competition among protocells^4–6^. The lipid world hypothesis, however, argues that both the encapsulated systems *and* the membrane compartment itself carry heritable information and contribute to protocellular fitness^7^ – a perspective that emphasizes the importance of the membrane and its composition. Indeed, simulations using the graded autocatalysis replication domain (GARD) model suggest that lipid assemblies can store and transmit their lipid compositions as heritable information, allowing specific compositions to adapt to environmental pressure^8–10^. Building upon these theoretical insights, compositional selection has been experimentally demonstrated with oil droplets^11^ and phospholipid-fatty acid mixed membranes^12–14^. Although oil droplets and fatty acid vesicles offer certain advantages in a prebiotic context due to their simplicity and plausibility^15^, a high permeability to biomolecules, protons, and other ions^16,17^, limits their ability to store compartmentalized metabolites and diminishes their relevance as effective models of a primordial cell. In contrast, phospholipid membranes exhibit an enhanced capacity for retaining compartmentalized contents^16–18^, and could be synthesized under prebiotic conditions^19–21^. Therefore, it is possible that protocells were made of phospholipids, while it is unclear if compositional selection could occur with these compartments.

As compartments generally grow in size before fission, the growth of a compartment is likely a prerequisite for the proliferation of protocellular systems^22^, and a fast growth rate within a certain environment would be a selective advantage for the protocell. While contemporary cells use sophisticated molecular machinery for compartment growth, primordial cells have only achieved growth through simpler mechanisms like vesicle fusion^3^. Among the various strategies for membrane fusion^23–31^, freeze-thaw cycles is a feasible driving force in a prebiotic context^32^ due to the universal potential for temperature cycling on early Earth^33^ and other planetary bodies like icy moons^34^. If phospholipid composition contributes to protocellular fitness, a selective enrichment of specific phospholipid compositions in the vesicle “offspring” should be detectable. Likewise, the encapsulated molecules, including metabolites and genetic information, should also be enriched even if they are selectively neutral. Yet, no experimental demonstration of these selection dynamics has been reported.

In this study, we experimentally explored phospholipid-dependent vesicle growth within an icy environment undergoing cycles of freezing and thawing (F/T). We found that vesicles with different lipid compositions exhibit variation in growth (Fig. 1A). When vesicles with different lipid compositions were mixed, F/T induced a selective partitioning of the phospholipid with more unsaturated bonds (Fig. 1B). Finally, we found that F/T cycling induced a selection for faster-growing phospholipid compositions that resulted in an enrichment of their compartmentalized genetic molecules, even though these materials were selectively neutral (Fig. 1C). This study is the first to demonstrate the selection of phospholipid composition and concurrent inheritance of compartmentalized genetic molecules in an icy environment, an environment that can trigger the growth of protocells.

**Fig 1.**
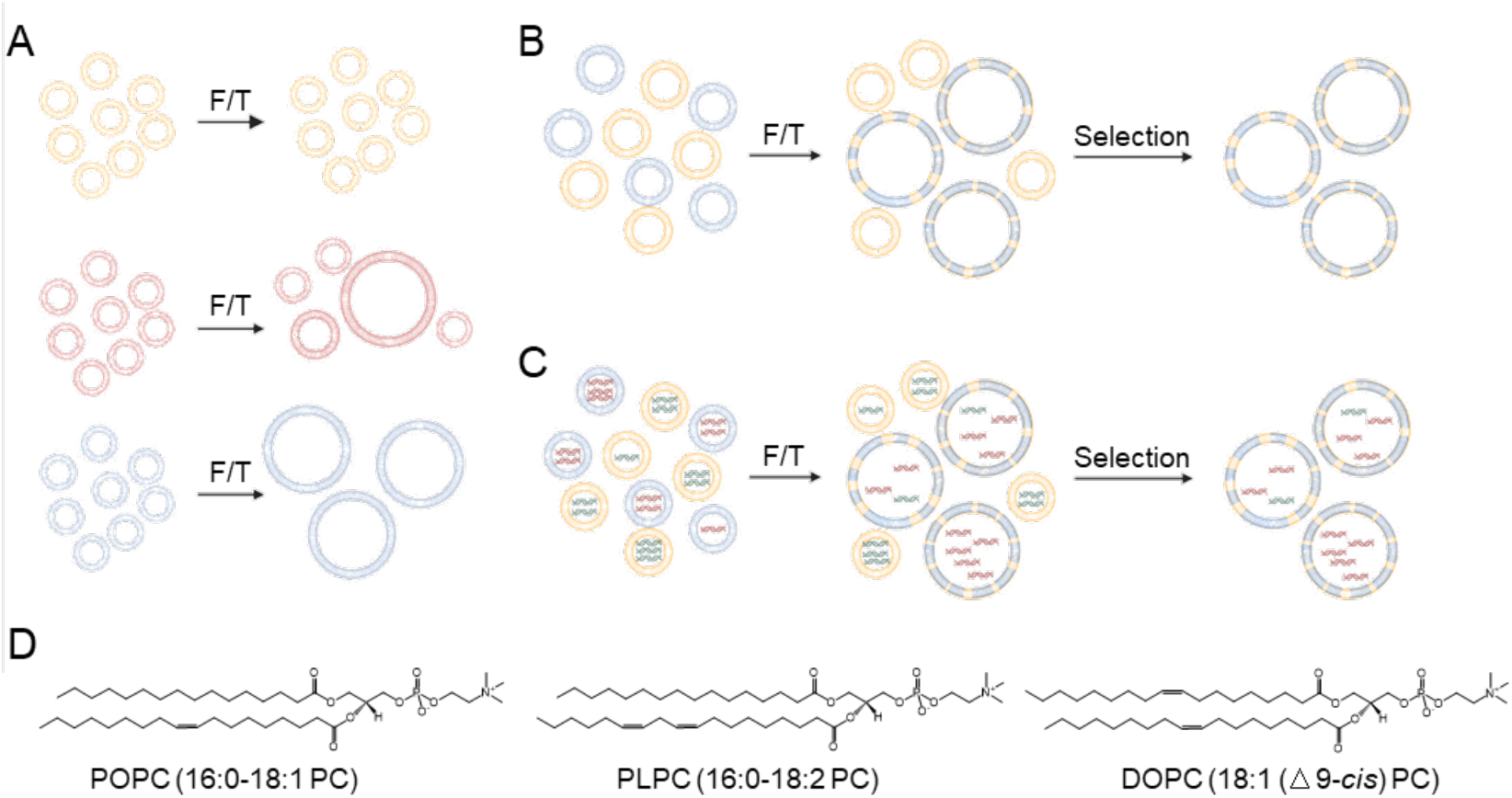
Schematics of freeze-thaw (F/T) cycles applied to vesicles composed of different phospholipids. (A) Examples of F/T with three different phospholipid compositions. Fusion occurs more efficiently in the order of blue, red, and orange vesicles. (B) F/T of a mixture of two different vesicles. The blue lipid is preferentially recruited into growing vesicles. (C) F/T of a mixture of two vesicles with different compositions of lipid. The encapsulated DNA is different but selectively neutral. (D) Chemical structures of the phospholipids used in this study: 1-palmitoyl-2-oleoyl-glycero-3-phosphocholine (POPC, 16:0-18:1 PC), 1-palmitoyl-2-linoleoyl-sn-glycero-3-phosphocholine (PLPC, 16:0-18:2 PC), 1,2-dioleoyl-sn-glycero-3-phosphocholine (DOPC, 18:1 (Δ9-Cis) PC).

## Results

### F/T cycles induce phospholipid-dependent vesicle fusion

We first investigated whether vesicles subjected to F/T conditions show variable degrees of growth depending on lipid compositions (Fig.1A). Large unilamellar vesicles (LUVs) with four different lipid compositions, *i*.*e*., i) 100% POPC, ii) 80% POPC: 20% PLPC, iii) 50% POPC: 50% PLPC, and iv) 100% PLPC, were prepared (Fig. 1D). The initial vesicle size (∼100 nm in diameter) was confirmed by dynamic light scattering (DLS) analysis (Fig. 2A, top). To increase fusion efficiency during F/T, vesicles were pelleted using an ultracentrifuge (630,000 g for 30 min at 4°C; hereafter, the term “pellet” refers to ultracentrifugation under these conditions), bringing the vesicles into proximity with each other. F/T was performed by freezing vesicles in liquid nitrogen for 1 min and then thawing at room temperature (∼24°C) for around 10 min until the ice completely melted. The thawed sample was vortexed prior to pelleting for subsequent F/T cycles (Fig. 2B). After repeating F/T three times (denoted as 3×F/T cycles hereafter), DLS measurements showed new peaks with vesicle diameters increase by one or more orders of magnitude compared to the original diameter for all phospholipid compositions tested (Fig. 2A, bottom). Note that peaks corresponding to fused growth were not observed for the samples without F/T or with F/T but without pelleting (Fig. S1).

**Fig. 2.**
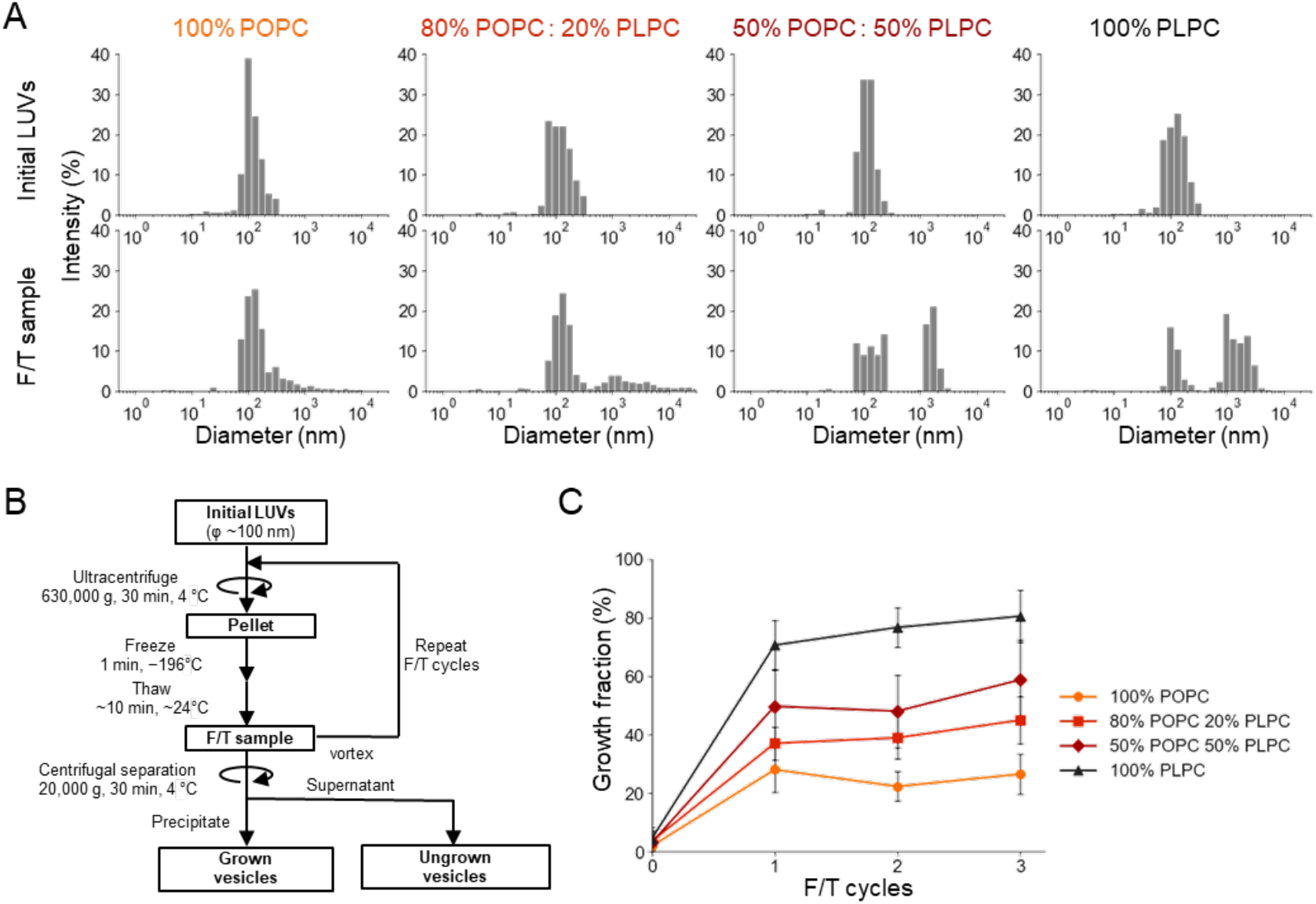
The F/T-induced growth of vesicles with four different initial phospholipid compositions. (A) Size distribution obtained using DLS measurements. The top panels show the results for initial LUVs while the bottom panels show the results after 3×F/T cycles. (B) Flowchart showing the F/T experimental procedure used in the present study. The ultracentrifuged LUV pellet was subjected to F/T followed by vortexing. Grown and ungrown vesicles were distinguished by centrifugal separation. (C) Growth fraction as a function of the number of F/T cycles. Each error bar represents the standard deviation calculated from three independent experiments (*n* = 3).

To assess the proportion of vesicles that grew, centrifugal separation at 20,000 g for 30 min at 4°C was performed on the vesicles after F/T cycles. The precipitated vesicles collected from this process were designated as “grown vesicles”, while “ungrown vesicles” refers to the vesicles that remained in the supernatant (Fig. 2B). We confirmed through DLS measurements that “ungrown vesicles” predominantly centered around their initial diameter of 100 nm, whereas “grown vesicles” showed a distinct shift toward larger sizes (Fig. S2). Using an enzymatic lipid quantification assay, we defined growth fraction as the mass fraction of lipids of grown vesicles over the total (sum of grown and ungrown vesicles). The near-zero values of growth fraction without F/T (0×F/T cycle in Fig. 2C) indicate that the centrifugal separation is mild enough to retain initial LUVs in the supernatant and does not remove the ungrown vesicles. In contrast, high growth fractions after 1–3×F/T cycles in all phospholipid compositions (Fig. 2C) were consistent with our DLS analyses (Fig. 2A). The growth fraction increased with the fraction of PLPC in the initial composition: Whereas 100% POPC (*i*.*e*., 0% PLPC) had a growth fraction of 27±7%, 100% PLPC (*i*.*e*., 0% POPC) had a growth fraction of 80±9% (Fig. 2C), with intermediate compositions have intermediate growth fractions. The one additional unsaturated bond in PLPC relative to POPC might have resulted in more destabilized lateral packing of the vesicle membrane, rendering vesicles with higher PLPC content more susceptible to fusion during F/T (see *Discussion* for details). In each case, the growth fraction is stable across successive rounds of F/T (Fig. 2C). When isolated ungrown vesicles were subjected to additional F/T cycling (Figs. S3 and S4), the growth efficiently was low, suggesting that these vesicles may be structurally distinct from vesicles that can grow. See Supplementary Information (Figs. S3 and S4) for details.

### PLPC increases membrane mixing efficiency during F/T-induced vesicle fusion

The grown vesicles after F/T cycles were sufficiently large for observation via laser scanning confocal microscopy (LSCM) (Fig. S5). When two LUV species with membranes containing fluorescently labeled lipids—one with nitrobenzoxadiazole (NBD)-tagged lipids and the other with rhodamine-tagged lipids—were mixed and subjected to 3×F/T cycles, the grown vesicles exhibited overlapping fluorescence signals (Fig. S5), indicating that membrane mixing had occurred in the grown vesicles. However, because of the lack of resolution of LSCM, it is not possible to quantify the mixing of membrane lipids at the molecular level. We thus used fluorescence resonance energy transfer (FRET) to quantitatively assess the membrane mixing mechanism (Fig. 3A). As membrane mixing occurs, the distance between the two fluorophores increases, leading to a decrease in the FRET signal and an increase in the NBD fluorescence (Fig.3A, right). Conversely, aggregation of LUVs results in little change in FRET signal (Fig.3A, left). We prepared LUVs containing both NBD- and rhodamine-tagged lipids as fluorescence donor and acceptor, respectively, and mixed the labeled LUVs with LUVs without any fluorophores (1:7 mass ratio) and subjected them to F/T cycles. We then measured the fluorescence of rhodamine and NBD, and calculated the FRET efficiency, *E*_*FRET*_, and used its reciprocal, 1/*E*_*FRET*_, as an indicator of the membrane mixing (Fig. 3A; see *Methods*); the larger the 1/*E*_*FRET*_ value is the more the membrane is mixed, and vice versa. We found that 1/*E*_*FRET*_ of grown vesicles increased after 1–3×F/T cycles compared to that of the initial LUVs (0×F/T cycle), irrespective of the initial phospholipid composition (Fig. S6A). Furthermore, 1/*E*_*FRET*_ after 3×F/T cycles increased along the PLPC percentage (Fig. S6A), indicating that a higher PLPC content resulted in grown vesicles with membranes better mixed than those with lower PLPC content.

**Fig. 3.**
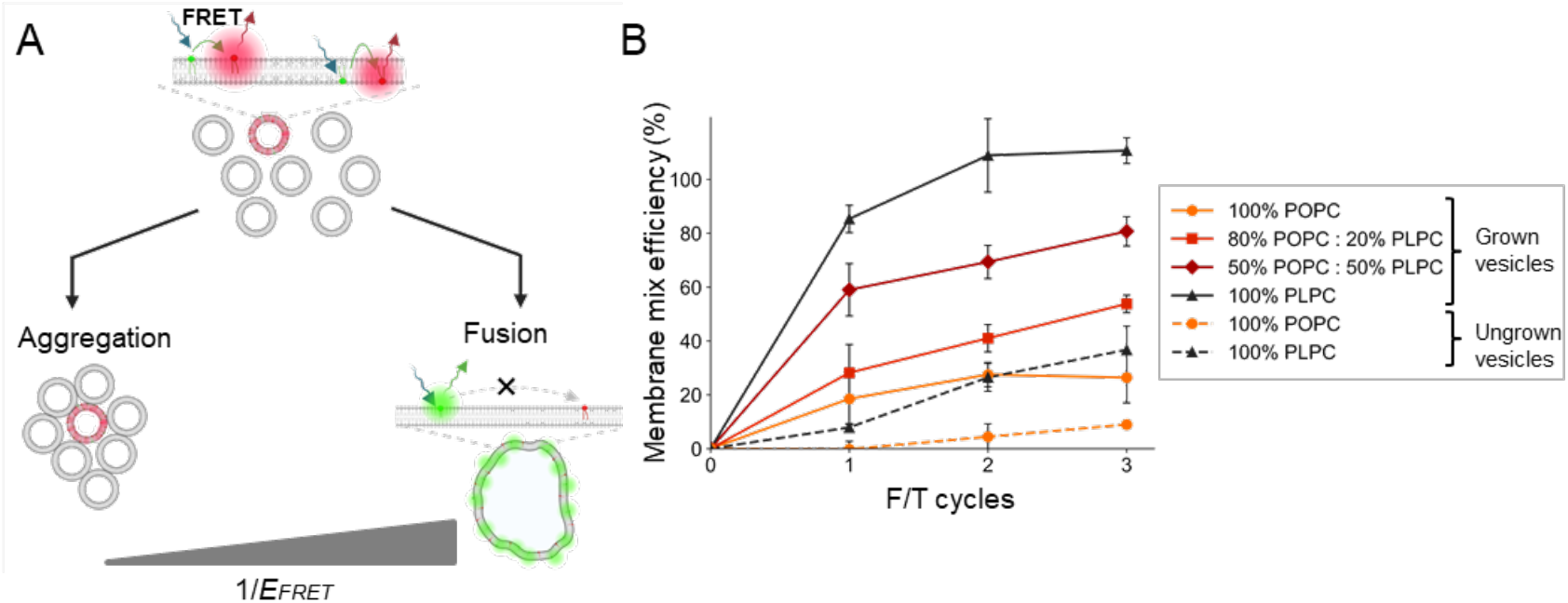
Membrane mixing of F/T-induced grown vesicles. (A) Schematic illustrating the FRET assay to quantify and distinguish two possible mechanisms for vesicle growth: vesicle aggregation (bottom left) and vesicle fusion (bottom right). The reciprocal of FRET efficiency, 1/*E*_*FRET*_, is expected to increase with the distance between the fluorescent donor, NBD (green dots), and the acceptor, rhodamine (red dots). (B) Membrane mix efficiencies of grown (solid lines) and ungrown (broken lines) vesicles after 1-3×F/T cycles.

We then converted 1/*E*_*FRET*_ into membrane mix efficiencies (see *Methods* for details, Fig. 3B). Here, 0% corresponds to aggregation of LUVs (*i*.*e*., no membrane mixing) (Fig. 3A, left) and 100% corresponds to homogenous mixing of lipids (Fig. 3A, right). After 2–3×F/T cycles, the membrane mix efficiency approached nearly 100% for 100% PLPC vesicles, while it remained below 30% for 100% POPC (*i*.*e*., 0% PLPC) vesicles. The first F/T cycle showed the most significant increase in the membrane mixing of grown vesicles compared to subsequent F/T cycles irrespective of phospholipid composition (Fig. 3B); this result mirrors that of the growth fraction (Fig. 2C), which also increased the most after the first F/T cycle. In contrast to grown vesicles, low membrane mix efficiencies (up to 30%) of ungrown vesicles (Fig. 2B) were observed (Fig. 3B, broken lines), which suggests a positive correlation between membrane mixing and vesicle growth.

Each error bar represents the standard deviation calculated from two independent experiments, each with three measurements (n = 6). 0% is defined as the 1/*E*_*FRET*_ value for the initial LUVs. 100% is defined as the 1/*E*_*FRET*_ value of LUVs containing both NBD-tagged and rhodamine-tagged lipids at a concentration assuming homogenous mixing of fluorescent: non-fluorescent LUV =1:7 (Fig. S6B).

### PLPC promotes mixing of encapsulated materials as a result of F/T-induced vesicle fusion

Vesicle fusion may also result in mixing or leakage of internal contents. To explore whether grown vesicles incorporate and maintain the contents compartmentalized within the initial LUVs that fuse to form the grown vesicles, two populations of LUVs encapsulating two different molecules, a calcein-cobalt (Co^2+^) complex and EDTA, respectively, were mixed at a 1:1 ratio and subjected to F/T cycles. (Fig. 4A). Upon content mixing, EDTA chelates Co^2+^, restoring calcein fluorescence that had been initially quenched by Co^2+^ (Fig. 4A), as observed by LSCM (Fig. 4B). We thus tracked calcein fluorescence across F/T cycles in the grown vesicles composed of four different phospholipid compositions (Fig. 4C). We confirmed that calcein fluorescence significantly increased after the first F/T cycle for all phospholipid compositions (Fig. 4C). After 3×F/T cycles, we saw a clear trend where the higher the PLPC percentage was, the lower the fluorescence intensity of the resulting grown vesicles (Fig. 4C).

**Fig. 4.**
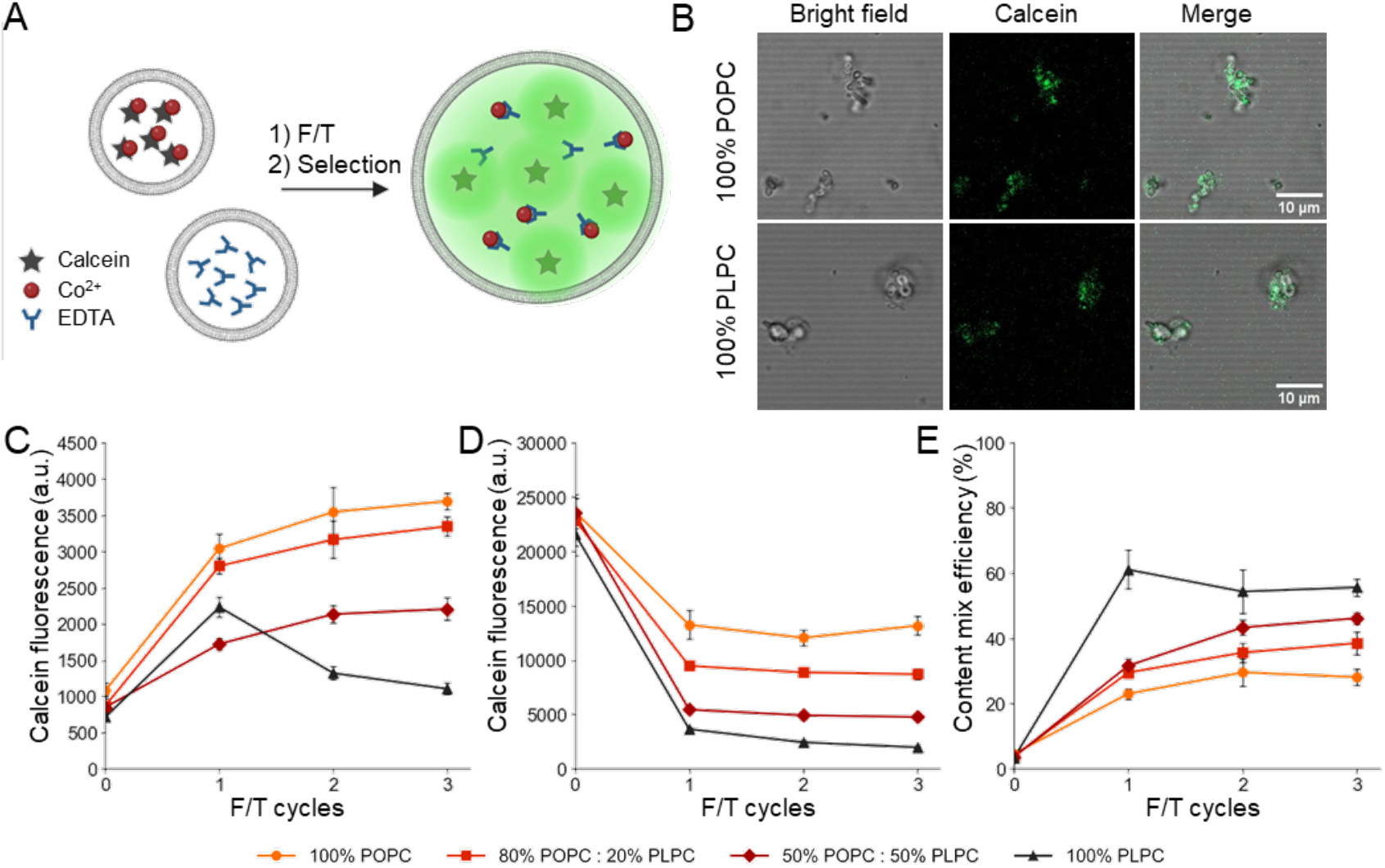
Content mixing and leakage of F/T-induced grown vesicles with different phospholipid compositions. (A) Schematic of the calcein-Co^2+^-EDTA assay for quantifying content mixing. Calcein exhibits fluorescence when Co^2+^ is removed by EDTA chelation. (B) Representative LSCM images of grown vesicles in the content mixing assay obtained by (i) bright-field observation, (ii) fluorescence observation with an excitation wavelength corresponding to calcein (488 nm), and (iii) a merged image of (i) and (ii). Imaging was performed using a 63×/1.40 oil-immersion objective lens at room temperature. (C) Calcein fluorescence in the initial LUVs (0×F/T cycle) and in grown vesicles after 1–3×F/T cycles. The fluorescence signals are derived from an equal amount of lipid vesicle (10 μg). (D) Calcein leakage over 1–3×F/T cycles. Vesicles used in this experiment were prepared with an internal content consisting of a mixture of the calcein-Co^2+^ complex and EDTA at the concentrations in (C). The fluorescence signals shown are derived from an equal amount of lipid vesicle (10 μg). Therefore, these data were used as the fluorescence value representing 100% content mix efficiency for each phospholipid composition at each F/T cycle. (E) Content mix efficiencies of LUVs over 1–3×F/T cycles calculated from data in (C) and (D). Each error bar represents the standard deviation calculated from two independent experiments, each with three measurements (n = 6).

Note that the fluorescence change observed in Fig. 4C is an overlay of i) content mixing and ii) content leakage triggered by F/T. To quantify the influence of content leakage on the change in the fluorescence observed in Fig. 4C, LUVs encapsulating fluorescent calcein made of four different phospholipid compositions were subjected to F/T cycles. Content leakage in the form of fluorescence intensity loss was observed in all LUV compositions tested and showed a clear dependency on phospholipid composition (Fig. 4D). The higher the PLPC content, the greater the fluorescence intensity decrease, which we speculate is likely due to the reduced membrane packing under F/T conditions. Finally, to extract the content mix efficiency of the LUV compositions tested (Fig. 4E), we normalized the calcein fluorescence intensity observed in grown vesicles (Fig. 4C) by subtracting the effect of content leakage (Fig. 4D, see *Methods*), and found that higher PLPC content resulted in higher content mix efficiency (Fig. 4E).

### Correlation between degree of lipid unsaturation and growth fraction, membrane mix efficiency, and content mixing efficiency

We have quantified the effect of phospholipid composition on the i) F/T-induced growth fraction (Fig. 2) and the ii) membrane mixing (Fig. 3) and iii) content mixing efficiencies (Fig. 4) of grown vesicles by varying the ratio of POPC and PLPC. Our results showed that a higher fraction of PLPC led to not only a higher growth fraction but also a more effective mixing of membranes and contents in the grown vesicles. Given that POPC has one double bond in the acyl chains and PLPC has two double bonds, we speculated that double bonds in the phospholipid promote fusion and thus growth, membrane mixing, and content mixing. We applied the same experimental setup to LUVs consisting of DOPC, a di-monounsaturated phospholipid, which has two double bonds in total, like PLPC, but in different acyl chains (Fig. 1D). Similarly to PLPC, 100% DOPC LUVs showed a higher growth fraction, membrane mix efficiency, and content mix efficiency than POPC LUVs over 1–3×F/T cycles (Fig. S7), indicating that one additional double bond, with relatively little dependence on its position, critically changes vesicle behavior.

We then introduced a parameter, “degree of unsaturation”, to compare our assay results across various compositions based on the number of double bonds in the lipid composition. Degree of unsaturation is taken to be an average number of double bonds in the acyl chains per phospholipid comprising the initial LUVs. We found a strong positive correlation (r > 0.9) between the degree of unsaturation and growth fraction (Fig. 5A). Furthermore, both the membrane and content mix efficiencies exhibited positive correlation (r > 0.9) with growth fraction (Figs. 5B and C), suggesting that F/T-induced grown vesicles can be selectively composed of growth-prone phospholipids, and simultaneously being enriched with contents originally encapsulated in growth-prone vesicles. Taken together, these data indicate that the degree of unsaturation reports on the ease of vesicle fusion, at least for the lipids used in these experiments.

**Fig. 5.**
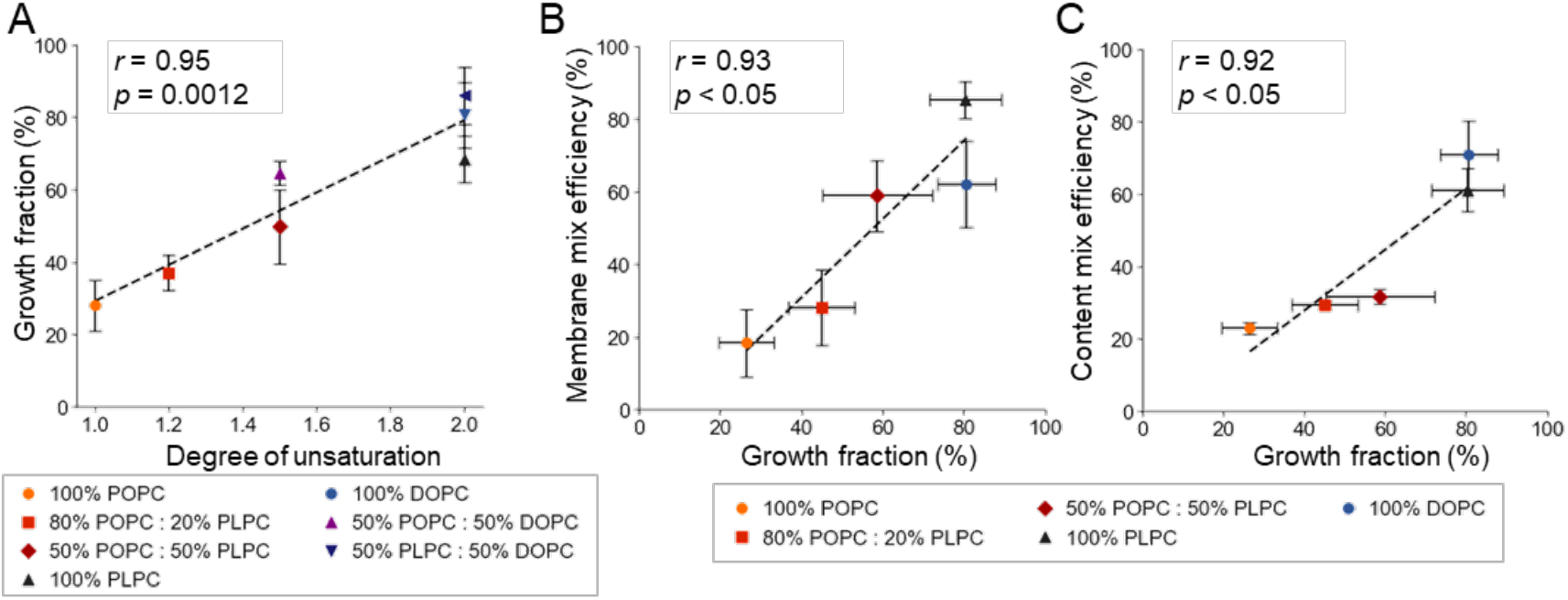
Comparison of vesicle behaviors after 1×F/T cycle among various phospholipid compositions. (A) Growth fraction as a function of “unsaturation level,” defined as the average number of double bonds in acyl chains per phospholipid constituting the initial LUVs. Correlation of growth fraction with (B) membrane mix efficiency and (C) content mix efficiency. Each data point corresponds to different phospholipid compositions of initial LUVs as indicated in the legend. Linear regression lines are provided as visual guides to illustrate the positive trend, along with Pearson correlation coefficients (*r*) and estimated *p*-values. Error bars represent standard deviations from three independent experiments (n = 3).

### F/T cycles induced the selection of a phospholipid and accompanied enrichment of genetic molecules

Assuming LUVs consisting of POPC and PLPC coexist, grown vesicles emerging as a result of the F/T cycles may selectively contain more PLPC than POPC due to the greater growth fraction of pure PLPC vesicles compared to pure POPC vesicles (Fig. 1B). In addition, as the content of PLPC vesicles tend to be transferred more to grown vesicles compared to POPC vesicles, the preference for PLPC may also lead to the enrichment of contents initially encapsulated in PLPC vesicles (Fig. 1C).

To test this hypothesis, we first developed an analytical method to quantify POPC and PLPC from mixed fractions using high-performance liquid chromatography with evaporative light scattering detection (HPLC/ELSD) (Fig. 6A and Fig. S8). We subjected a 1:1 weight mixture of 100% POPC LUVs and 100% PLPC LUVs to F/T cycles. As expected, the PLPC/POPC ratio of the grown vesicles increased approximately two-fold (0.98±0.07 to 1.83±0.13) after 3×F/T cycles, indicating selective incorporation of PLPC into the grown vesicles (Fig. 6B). We also investigated whether encapsulated molecules in the initial vesicle population were enriched by using DNA as a model analyte (Fig. 1C). DNA encoding green fluorescent protein (GFP-DNA, 956 bp) and mScarlet protein (mScarlet-DNA 929 bp) were encapsulated in 100% POPC LUVs and 100% PLPC LUVs, respectively. These two LUV populations were mixed at a 1:1 lipid mass ratio and subjected to 1–3×F/T cycles. DNAs encapsulated in the initial LUV mixture and the grown vesicles were recovered and subjected to quantitative PCR (qPCR). As a result, following each F/T cycle (Fig. 6C), the copy number ratio of mScarlet-DNA to GFP-DNA increased by two-fold (1.96±0.15 to 4.31±0.08) after 3×F/T cycles (Fig. 6D). We find that grown vesicles that form through F/T cycles exhibited both a bias toward PLPC *and* preferential retention of DNA, while selectively neutral, originating from PLPC LUVs. These results demonstrate that the enrichment of compartmentalized genetic molecules in the absence of selection pressure can be driven by F/T-induced selection of membrane lipids.

**Fig. 6.**
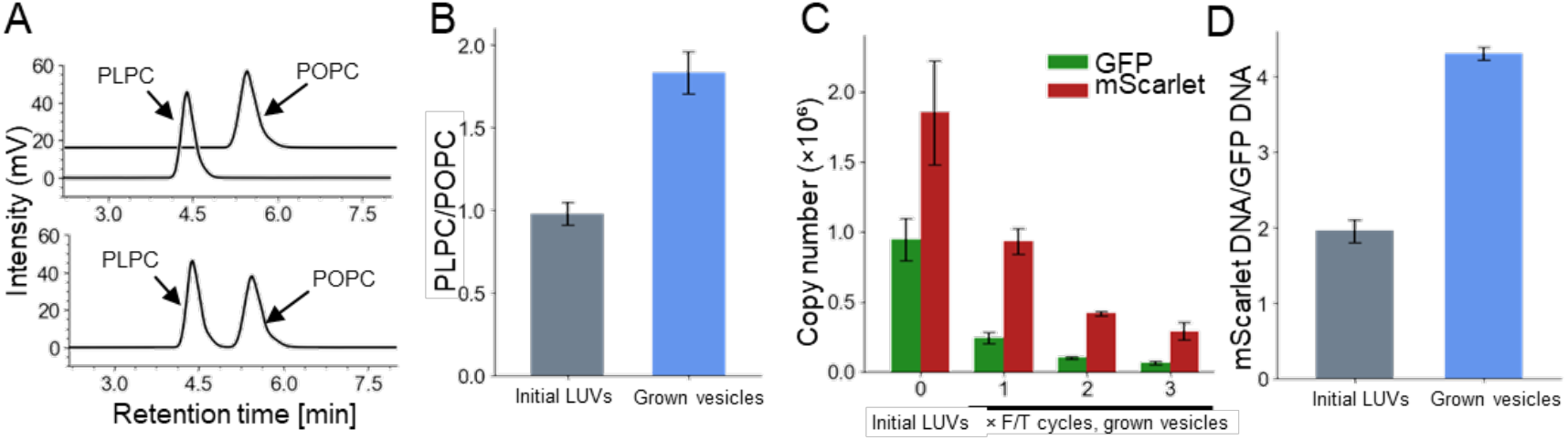
Selection of membrane lipids and the coupled enrichment of encapsulated genetic molecules. (A) Representative chromatogram from HPLC/ELSD analysis of POPC and PLPC, measured separately (top) and as a 1:1 mass ratio mixture (bottom) (see *Methods* and Fig. S8 for details). (B) Changes in the PLPC/POPC mass ratio over F/T cycles. Data from the initial LUVs and the grown vesicles collected after 3×F/T cycles are compared. (C) Copy numbers of GFP-DNA (green) and mScarlet-DNA (red) extracted from the initial LUV mixture and the grown vesicles obtained after 1–3×F/T cycles. DNA was quantified by qPCR. (D) The change in mScarlet-DNA/GFP-DNA molar ratio from the initial LUVs compared to that from the grown vesicles after 3×F/T cycles. Error bars represent the standard deviation from three independent experiments (n = 3).

## Discussion

Here, we demonstrated that phospholipid composition contributes to vesicle growth under F/T cycling conditions. We compared three types of phospholipids—POPC, PLPC, and DOPC—which share the same head group but differ in the number and position of double bonds in their acyl chains. Higher PLPC and DOPC content in initial vesicles, rather than higher POPC content, resulted in greater growth fractions, membrane mix efficiency, and content mix efficiency (Figs. 2–5), strongly suggestive of a fusion-based growth mechanism. The positive correlation among these three properties further suggests that growth-prone vesicles can effectively supply growth-prone lipids and encapsulated contents to the resulting population of grown vesicles, preserving informational continuity between generations of vesicles (Fig. 6).

As membrane components, we used phospholipids, in particular phosphatidylcholine (PC), owing to their structural and chemical continuity with those of contemporary cells, and their ability to be synthesized under prebiotic condition^19–21^ and to retain compartmentalized essential contents. In particular, our experimental results comparing three PC lipids highlighted that only a small difference in the acyl chain is sufficient to dramatically change the membrane behavior under F/T conditions, leading to the selection of lipids and the concurrent enrichment of encapsulated content. While further experiments are needed, we speculate that similar selection could occur among other amphiphilic molecules with greater chemical and structural differences, including phospholipids other than PC as well as fatty acids, which could have been more easily synthesized on early Earth than di-acyl chain phospholipids^1,35^.

We speculate that the superior growth of PLPC and DOPC vesicles compared to POPC under F/T cycles can be attributed to differences in their physicochemical properties, such as phase transition temperatures (T_m_) or membrane lateral packing properties. The T_m_ values of POPC, PLPC, and DOPC are -2°C, -18°C, and -17°C, respectively^36^. Given that vesicle fusion can occur in confined spaces between ice crystals at subzero temperatures, vesicles composed of lipids with lower T_m_, such as PLPC and DOPC may retain higher membrane fluidity in these confined spaces, facilitating fusion. However, in our experimental setup, vesicle confinement was already ensured by ultracentrifugation, making additional ice crystal-induced confinement unlikely to play a major role. The limited influence of ice crystal confinement is further supported by the observation that the growth fraction generally exhibited small differences depending on whether the frozen sample was thawed moderately at 24°C or rapidly at 65°C across the three phospholipids over 1–3×F/T cycles (Fig. S9). Therefore, under the conditions where vesicles were brought into close contact via ultracentrifugation, we suppose that membrane lateral packing, rather than T_m_ of individual phospholipids, plays a more significant role in vesicle growth. Compared to POPC, more unsaturated phospholipids like DOPC and PLPC tend to form less densely packed membranes^37,38^. Under the stresses of ice crystal formation, membranes can become destabilized or fragmented, requiring structural reorganization upon thawing. In the case of PLPC and DOPC, their loosely packed lateral organization may expose more hydrophobic regions during membrane reconstruction, facilitating interactions with adjacent vesicles and making fusion energetically favorable.

F/T cycles in nature, driven by geological or environmental processes such as diurnal and seasonal temperature changes, or tidal-induced ice-ocean convections^34,39,40^ occur much more slowly than our experimental timescales. Such slow change in temperature can lead to the gradual growth/dissolution of ice crystals, which in turn exclude/retain solutes within the inter-crystal liquid phase, resulting in eutectic concentration. It is plausible, then that vesicles accumulate in the confined eutectic phase, and thus, fusion processes could have occurred within this locale. We employed ultracentrifugation, which does not exist in the natural setting, to efficiently induce vesicle-vesicle contact before freezing. We nevertheless consider ultracentrifugation as an analogous condition to the eutectic concentration because both play a role in concentrating the vesicles in a confined space.

As a plausible prebiotic driver for chemical evolution, dry-wet cycles have been well discussed^41,42^. However, the drying conditions may cause excessive dehydration and heat stress, leading to irreversible denaturation of molecules including proteins and lipid bilayer. We believe that F/T cycles can be an effective environment when the reaction and the molecules require water molecules. Indeed, F/T conditions have been reported to be conducive to key prebiotic pathways, such as synthesis of RNA bases^43^, nonenzymatic template-directed polymerization of RNA^44^, ligation of RNA oligomers forming functional ribozymes^45^, and effective hybridization of kilobase-sized DNA fragments^46^. F/T cycles also induce the supply of encapsulated molecules into vesicles from other vesicles ^32,47–49^. Here, we add to the prebiotic repertoire of F/T by demonstrating how F/T cycles can directly effect vesicle behavior depending on physicochemical properties of lipid membranes. Our results, together with those of others suggest that icy conditions could have played a significant role in facilitating the selective assembly of biomolecules and molecular systems.

We showed that genetic information can be enriched and inherited within the grown vesicles based on their lipid composition. When assuming the presence of compartments harboring various molecules, including diverse genetic information, our observation suggests that the icy environment may contribute to mixing and *inheriting* mixed genetic information to the offspring based on the chemical composition of the compartment. Once the encapsulated system has inherited and gained sufficient materials, the intra-vesicular system might start contributing more to the protocellular fitness than the membrane composition, a step forward to contemporary cellular systems. Finally, by integrating F/T cycle-driven selection of grown vesicles with vesicle fission mechanisms, a recursive selection of vesicular systems across successive generations may be realized. Such fission could be achieved through means such as osmotic pressure changes or mechanical shear, consequently leading to the emergence of a primordial cell capable of Darwinian evolution.

## Methods

### Chemicals

1-palmitoyl-2-oleoyl-glycero-3-phosphocholine (POPC), and 1,2-dioleoyl-sn-glycero-3-phosphocholine (DOPC) were purchased from NOF Corp. 1-palmitoyl-2-linoleoyl-sn-glycero-3-phosphocholine (PLPC), 1,2-dioleoyl-sn-glycero-3-phosphoethanolamine-N-(lissamine rhodamine B sulfonyl)(ammonium salt) (18:1 Liss Rhod PE) and 1,2-dioleoyl-sn-glycero-3-phosphoethanolamine-N-(7-nitro-2-1,3-benzoxadiazol-4-yl) (ammonium salt) (18:1 NBD PE) were purchased from Avanti Polar Lipids Inc. Calcein, cobalt(II) chloride hexahydrate, magnesium chloride hexahydrate, and calcium chloride dihydrate were purchased from FUJIFILM Wako Pure Chemical Corp. Ethylenediaminetetraacetic acid disodium salt (EDTA) was purchased from Tokyo Chemical Industry Co., Ltd. Sodium chloride, disodium hydrogenphosphate dodecahydrate, potassium chloride, and potassium dihydrogenphosphate, used for preparing PBS buffer, were purchased from Nacalai Tesque, Inc. Triton X-100 was purchased from Kishida Chemical Co., Ltd. Plasmid DNAs encoding a fluorescent protein were constructed in the previous study^50^. Briefly, green fluorescent protein (frGFP) gene^51^ was amplified from pUC-frGFP vector (Musaiboukun, Taiyo Nippon Sanso) and sub-cloned into pIVEX2.3 vector using HiFi DNA assembly kit (New England Biolabs). Red fluorescent protein (mScarlet-I3) gene^52^ was synthesized in Eurofins Genomics, and sub-cloned into pIVEX2.3 vector. The DNA sequences were verified by Sanger sequencing. The DNAs used in this experiment were PCR products amplified from these plasmids using primers T7F (5’-ATGCGTCCGGCGTAGAGGATCGAGA) and T7R (5’-AGGGGTTATGCTAGTTATTGCT CAGCGG), and KOD polymerase (Toyobo) according to manufactures instruction.

### Preparation of large unilamellar vesicles (LUVs)

∼100 nm size large unilamellar vesicles (LUVs) were prepared by extrusion methodology^53^ with some modification. Briefly, 10 mg of lipid dissolved in ∼200 μl of pure chloroform was rotated and evaporated under a vacuum in a round-bottom flask for 1 h. The resulting dried lipid film was hydrated with 300 μl of PBS buffer (137 mM NaCl, 8.1 mM Na_2_HPO_4_, 2.68 mM KCl, 1.47 mM KH_2_PO_4_, pH 7.4) unless specified, to yield 300 μl of ∼30 mg/mL vesicle solution (multilamellar vesicles). The vesicle solution was sonicated (VS-1003, AS ONE Corp.) for 10 min (repeated for a total of 10 min: 28 Hz for 1min, 45 Hz for 1min, 100 Hz for 3s) and vortexed for 15 s to resuspend completely. The suspension was then subjected to 5 cycles of repeated F/T (incubated in liquid nitrogen for 1 min, then incubated in 65°C of water bath until thawed). The vesicle suspension was extruded 10 times using an extruder (Avanti Polar Lipids) through a 100 nm polycarbonate membrane filter. Subsequently, centrifugal separation at 20,000 g for 20 min was conducted to eliminate aggregates.

### F/T procedure for vesicle fusion

500 μl of LUV solution was ultracentrifuged at 630,000 g for 30 min at 4°C. The supernatant was removed, and the precipitated fraction was frozen in liquid nitrogen at -196°C for 1 min. The frozen precipitation was then thawed for ∼10 min at room temperature of approximately 24°C. After the pellet was thawed, it was resuspended in 600 μl of PBS buffer and 100 μl of the solution was collected for analysis. The remaining 500 μl was subjected to the next round of F/T cycles.

### Characterization of LUVs

The size distribution of vesicles before and after F/T cycles was determined by dynamic light scattering analysis (DelsaMax CORE, Beckman Coulter, Inc). Measurement was conducted with a laser wavelength was 660.9 nm, and the temperature in the instrument was kept at 25°C. For each sample, three measurements of 10 runs were performed, and the histogram was generated using the average DLS intensity across these three measurements. The mass of phospholipids was measured using LabAssay™ Phospholipid kit (Choline Oxidase DAOS method, FUJIFILM Wako Pure Chemical Corp.) according to manufacturers’ instruction. Images of vesicles were acquired using a confocal laser scanning microscope LSM 900 (Carl Zeiss), equipped with a 63×/1.40 oil immersion objective at room temperature. The fluorescence imaging was performed with an excitation wavelength of 561 nm for Rhodamine, 488 nm for NBD, and 488 nm for calcein.

### Quantitative analysis of membrane mixing of vesicles using fluorescence resonance energy transfer (FRET)

Three sets of LUVs were prepared: (i) fluorescently labeled vesicles containing 0.2 mol% 18:1 NBD-phosphatidylethanolamine (PE) and 0.2 mol% 18:1 Liss Rhod-PE, (ii) non-fluorescent labeled vesicles (*i*.*e*., without fluorescent lipids), and (iii) fluorescently labeled vesicles containing 0.025 mol% 18:1 NBD-PE and 0.025 mol% 18:1 Liss Rhod PE. For the membrane mixing experiments, (i) and (ii) vesicles were mixed at a 1:7 (w/w) ratio to prepare a 500 μL vesicle solution with a total lipid concentration of 2.67–3.33 mg lipid/mL. This mixture was subjected to F/T cycles, and the collected sample was adjusted to a concentration of 0.15 mg lipid/mL for analysis. Fluorescence measurements were taken using a multi-mode microplate reader (Bio Tek Instruments, Inc.) FRET rhodamine fluorescence by Ex: 460 nm/Em: 595 nm (*I*_*Rhod*_) and NBD fluorescence by Ex: 460 nm/Em: 540 nm (*I*_*NBD*_) were measured. The relative FRET efficiency was defined as follows: *E*_*FRET*_ = *I*_*Rhod*_/*I*_*NBD*_. The inverse *E*_*FRET*_ (1 / *E*_*FRET*_) served as a measure of membrane mixing. As a control to obtain the 1/*E*_*FRET*_ for the fully membrane-mixed state, the data derived from sample (iii) was used. To quantify membrane mixing efficiency, the 1 / *E*_*FRET*_ value of the initial LUVs (0×F/T cycle) was set as 0%, and the 1 / *E*_*FRET*_ value obtained from sample (iii), representing complete lipid mixing, was set as 100%. Membrane mixing efficiencies for all conditions were then calculated by normalizing the observed 1 / *E*_*FRET*_ values between these two references.

### Quantitative analysis of content mixing of vesicles using calcein-Co^2+^-EDTA assay

Three distinct sets of vesicles were prepared: (i) vesicles loaded with 0.1 mM calcein and 0.2 mM CoCl_2_, (ii) vesicles loaded with 2.5 mM EDTA, and (iii) vesicles loaded with 0.05 mM calcein, 0.1 mM CoCl_2_, and 1.25 mM EDTA. These vesicles were prepared in PBS buffer containing the final concentrations of calcein, CoCl_2_, or EDTA as described above. Ultracentrifugation was performed to remove the unencapsulated chemicals and to replace the external solution with PBS buffer. For the content mixing experiments, vesicles (i) and (ii) were mixed at the ratio of 1:1 (w/w) at a final lipid concentration of 2.67 mg lipid/ml and the F/T was performed, with each cycle followed by washing the vesicle solution three times by ultracentrifugation at 630,000 g for 30 min at 4°C, to ensure the removal of calcein outside the vesicles. The collected sample was adjusted to a concentration of 0.20 mg lipid/mL for further analysis. Calcein fluorescence intensity was measured using a multi-mode microplate reader (Ex: 490 nm, Em: 520 nm). As a control for the fully content-mixed state, the values obtained using vesicle (iii) were used. Calcein leakage was quantified by measuring the decrease in fluorescence intensity of vesicle (iii) after each F/T cycle. To normalize the fluorescence intensity into content mix efficiency, the intensity of vesicle mixture of (i) and (ii) was divided by that of vesicle (iii) after subtracting the background intensity from respective results.

### Quantitative analysis of the lipid composition within phospholipid vesicles

The phospholipid composition of the vesicles was analyzed and quantified using a High-Performance Liquid Chromatography system (Prominence Ultra-Fast Liquid Chromatography, Shimadzu Corp.) with an Evaporative Light Scattering Detector (ELSD-LT II, Shimadzu Corp.). Prior to the chromatographic analysis, the vesicles were disrupted using 0.1% (v/v) Triton X-100 to ensure homogeneity. A volume of 5 µL of the resulting lipid mixture was injected. The separation was carried out using an XBridge C8 column (2.5 µm, 4.6 × 300 mm, Waters) at 55°C, where the mobile phase consisted of 0.1% (v/v) formic acid in methanol-water (84:16, v/v) over a 10 min run at a flow rate of 1.0 mL/min. Detection was achieved using the ELSD with a drift tube temperature set at 40 °C, a gain of 2, and a nitrogen (N_2_) pressure of 340 kPa.

### Quantitative analysis of the copy number of DNA within phospholipid vesicles

Two distinct populations of vesicles were prepared in PBS buffer containing 2.5 mM MgCl_2_ and 0.5 mM CaCl_2_: (i) 100% POPC vesicles hydrated with 10 nM DNA encoding green fluorescent protein (GFP) and (ii) 100% PLPC vesicles hydrated with 10 nM DNA encoding mScarlet. To remove external DNA, each vesicle population underwent DNase I (Takara Bio Inc.) treatment. Following DNase I digestion, the samples were incubated at 75°C for 10 min to inactivate the enzyme. The two LUV populations were mixed at a 1:1 (w/w) ratio, and F/T cycles were performed. The collected samples were adjusted to a concentration of 0.20 mg lipid/mL, and DNase I treatment was conducted to remove any DNA that had leaked into the external solution, followed by the inactivation of enzyme through incubating at 75°C for 10 min. Subsequently, 0.5% (v/v) Triton X-100 was added to disrupt the vesicle membranes. The DNA that was inside the vesicles was then purified (Nucleospin® Gel and PCR Clean-up, Takara Bio Inc.) according to the manufacturer’s instructions. Quantitative PCR (qPCR) was performed with TB Green^®^ *Premix Ex Taq*™ II kit (Takara Bio Inc.), using a primer set (CGCCATAACATCGAAGACGG, AGGCTGATTGCGTAGACAGG) to detect GFP, and a primer set (CAAGACCACCTACAAGGCCA, GCGTTCGTACTGTTCCACCA) to detect mScarlet gene.

## Acknowledgments

This study was supported by the Human Frontier Science Program Grant Number RGP003/2023 (TM), JSPS KAKENHI Grant Numbers 22KJ1296 (NN), 22K21344 (TM), and 21H05228 (TM), and the Astrobiology Center Program of the National Institutes of Natural Sciences (NINS) Grant Number JY230122nn (TM).

## Author Contributions

TS performed most of the experiments, YT contributed to sample analysis, TW established the experimental protocol for ELSD measurement. TS, KK, YS, TM conceived the project, TS, NN and TM wrote the paper, and all authors discussed the project and edited the manuscript. We thank Tony Z. Jia and Liam M. Longo (ELSI, Institute of Science Tokyo) for the critical reading of the manuscript and many constructive comments, and Prof. Yoshikazu Tanaka (Tohoku University) for TEM analysis of the vesicles at the early phase of the project.

## Notes

Competing interests

The authors declare no competing interests.

## Notes

### Competing Interest Statement

The authors have declared no competing interest.

### Summary of Updates

Mistake was found in the author list and the mansucript

