## Supplementart Information for "Compositional selection of phospholipid compartments in icy environments drives the inheritance of encapsulated genetic information"

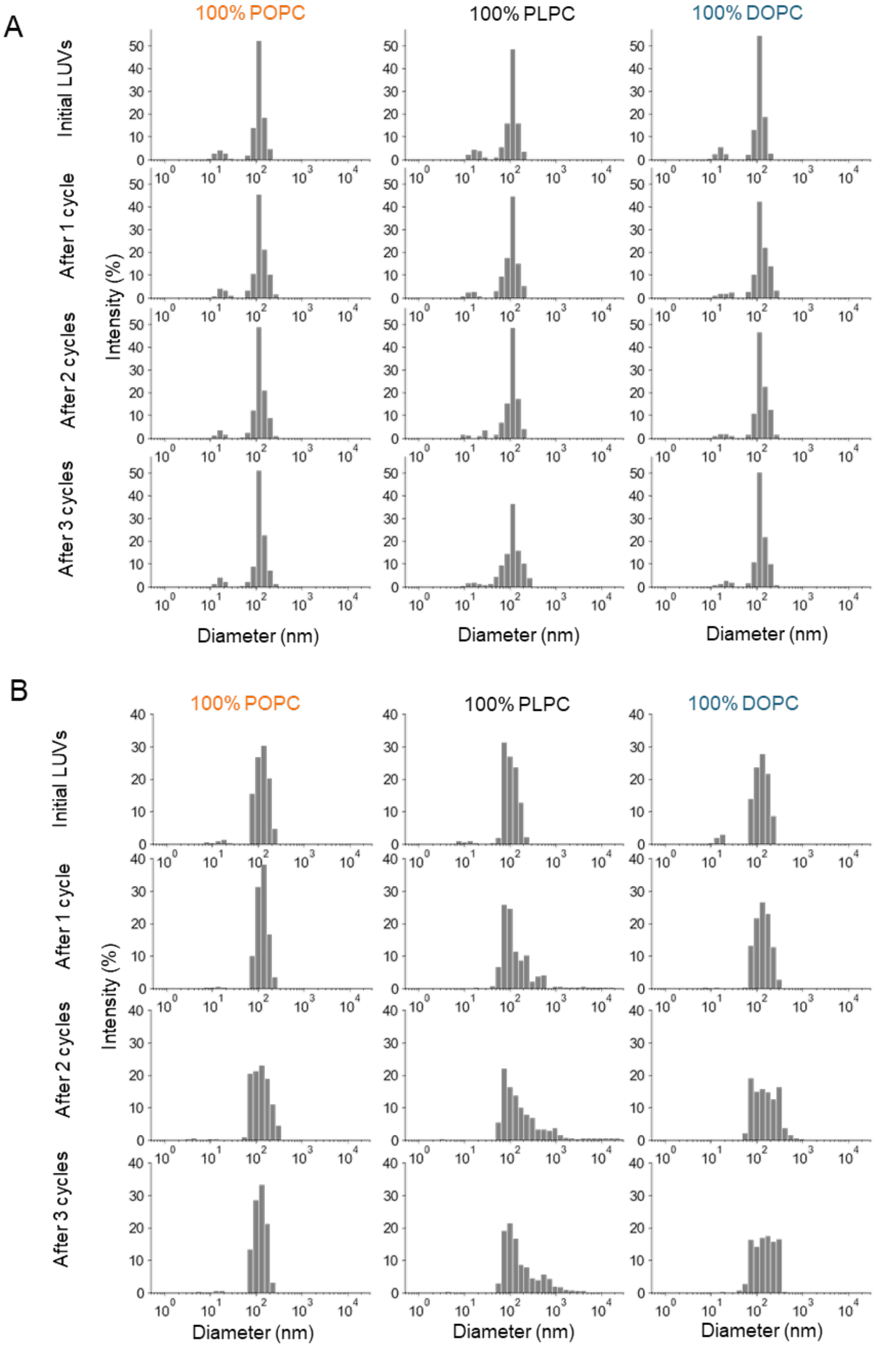


**Figure S1**: The effect of pelleting and F/T on the size of the vesicles. (A) The LUVs at 1.5 mg lipid/mL in PBS were subjected to repeated pelleting using ultracentrifugation and vortexing for resuspension. Size distribution was obtained using DLS measurement. The histogram shows that the size distribution is not affected by pelleting and vortexing alone. (B) The LUVs at 1.5 mg lipid/mL in PBS were subjected to F/T cycles as described in the Method section, except that LUVs were not pelleted before freezing. The peaks in the size distribution one order of magnitude or larger than the initial LUVs were rarely observed, indicating the necessity of pelleting for efficient vesicle growth by F/T cycles.

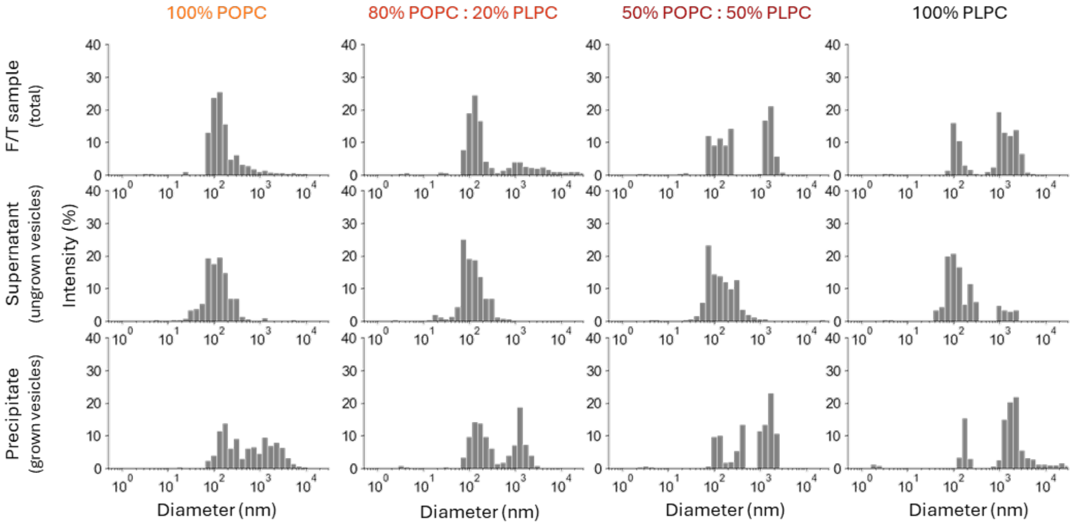


**Figure S2**: DLS analyses data after 3×F/T cycles. The preparation method for F/T sample (top), Supernatant (middle), and Precipitate (bottom) are shown in Fig. 2B. Large particles appeared after 3×F/T cycles (total; identical to the data shown in Fig 2A, lower panel). After centrifugation at 20,000 g, 30 min at 4ºC, the supernatant and precipitate were analyzed by DLS measurement. Large particles were enriched in the precipitate.

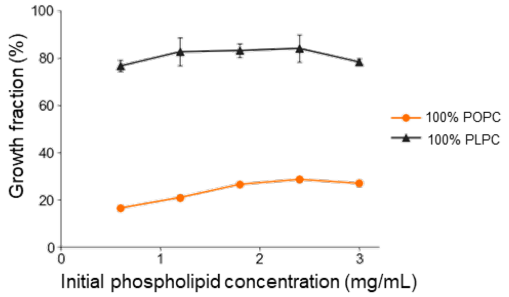


**Figure S3:** The effect of lipid concentration on the growth fraction of LUVs after 1×F/T cycle. We observed a notable increase in growth fraction after the first F/T step, but little increase in growth fraction after further F/T cycles (Fig. 2C). One possible explanation is that the upper limit of growth fraction is determined by the phospholipid concentration; however, this was inconsistent with the observation that the initial concentration of the vesicle hardly affected the growth fraction (Fig. S3). Here, the experiments were done with 100% POPC and 100% PLPC vesicles. The growth fraction was hardly affected by the initial phospholipid concentrations between 0.5–3 mg/mL. The F/T cycle and the quantification procedures are the same as described in the main text. While the data are with 100% POPC and 100% PLPC vesicles, 80% POPC: 20% PLPC and 50% POPC: 50% PLPC vesicles are likely to exhibit the same lipid concentration dependency. Each error bar shows the standard deviation ($n = 3$).

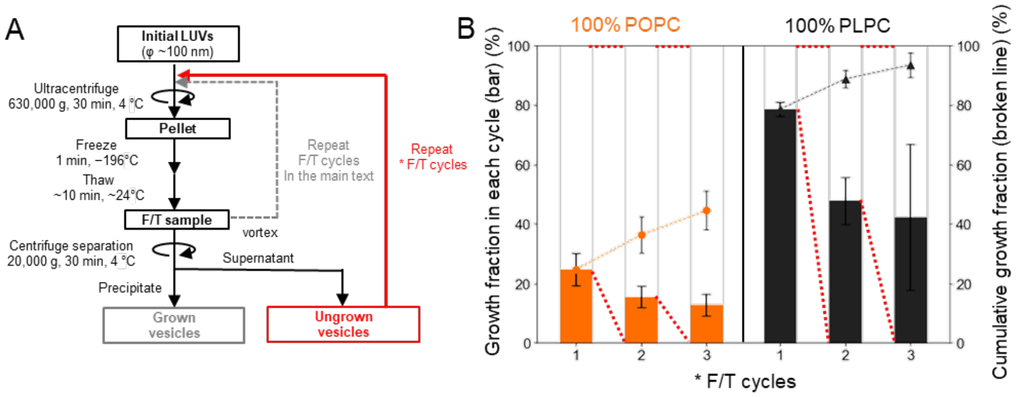


**Figure S4**: The effect of F/T cycles on ungrown vesicles. We observed a notable increase in growth fraction after the first F/T step, but little increase in growth fraction after further F/T cycles (Fig. 2C). One possibility is that the ungrown vesicles have low fusion efficiency. Indeed, the growth fraction was low when the isolated ungrown vesicles were subjected to an additional F/T cycle (Fig. S4). These observations suggest that even a single F/T cycle can alter the vesicle growth behavior. (A) Flowchart of the F/T cycles of an ungrown vesicle, indicated in red (* F/T cycles). The ungrown vesicle obtained as a supernatant after centrifugation was subjected to the following F/T cycle. (B) The growth fraction in each cycle, quantified in the same way as the main text (bars), was relatively lower in the 2nd and 3rd * F/T cycles than the 1st F/T cycle for both 100% POPC and 100% PLPC, despite that phospholipid concentration has little effect on the growth fraction (Fig. S3). When the cumulative growth fraction (broken lines) was plotted based on the growth fraction in each cycle, a similar deceleration in growth after the initial cycle, as observed in Fig. 2C, was confirmed. This result suggests that the stagnation of the growth fraction after the initial cycle may be attributable to the relatively low fusion efficiency of vesicles that remained ungrown.

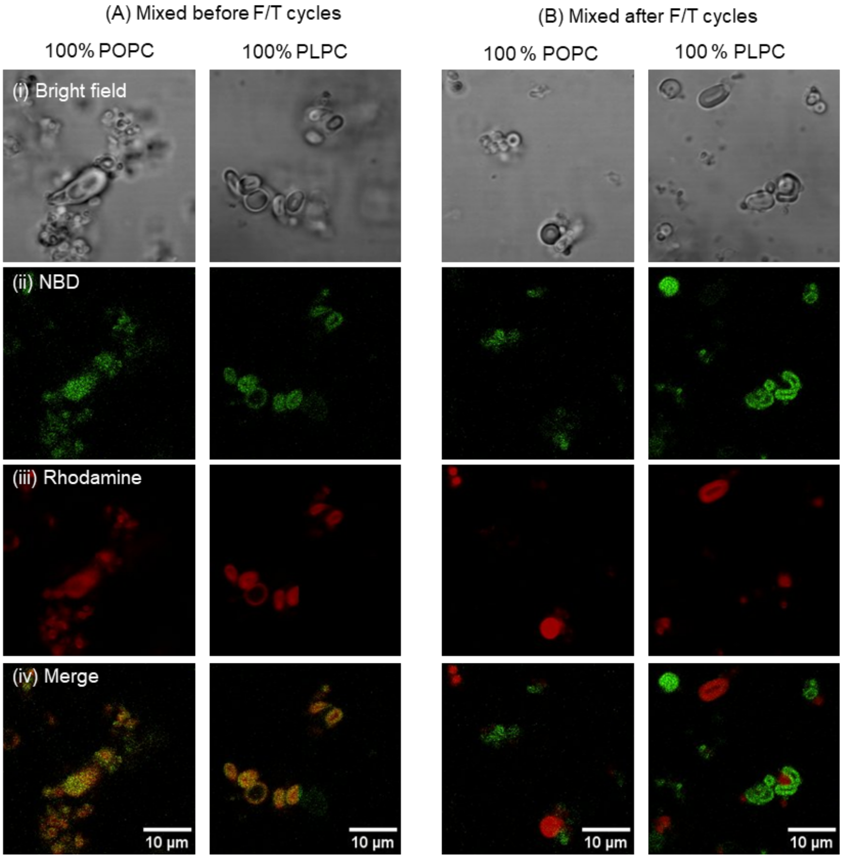


**Figure S5**: Laser scanning confocal microscopy (LSCM; LSM 900, Carl Zeiss) images of the grown vesicles by (i) bright field, (ii) rhodamine (ex. 561 nm), (iii) NBD (ex. 488 nm), and (iv) the merged of (ii) and (iii). Imaging was performed with a 63×/1.40 oil-immersion objective lens at room temperature. The initial LUVs were fluorescently labeled by either NBD or rhodamine lipid at 0.2 mol%. These two populations of vesicles were mixed in 1:1 lipid mass ratio and subjected to 3×F/T cycles ((A) mixed before). As a control (B) 3 × F/T was subjected to NBD or rhodamine containing LUVs respectively and mixed ((B) mixed after). (A) shows the overlayed fluorescence of NBD and rhodamine, while (B) does not. These show that the LSCM images can be used to trace F/T-induced membrane mixing of both 100% POPC and 100% PLPC LUVs.

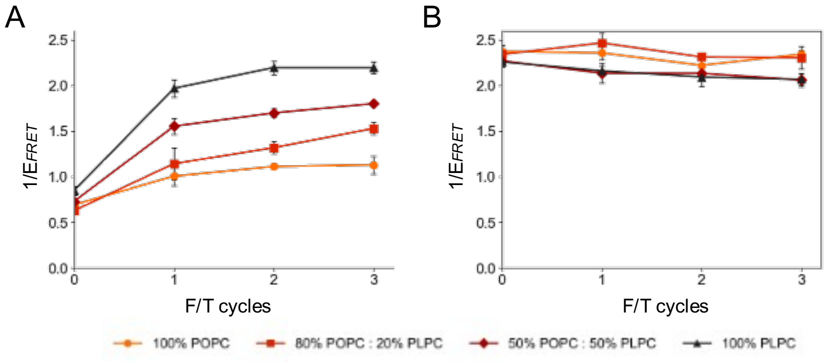


**Figure S6**: Membrane mixing assay. (A)The results of $1/E_{FRET}$ for the initial LUVs (0×F/T cycle) and the grown vesicles after 1–3×F/T cycles. The fluorescently labeled LUVs (containing both NBD-tagged and rhodamine-tagged lipids as fluorescence donors and acceptors, respectively) were mixed with non-fluorescent LUVs at 1:7 mass ratio) and subjected to F/T cycles. (B) The $1/E_{FRET}$ values derived from the F/T cycles of LUVs containing both NBD-tagged and rhodamine-tagged lipids at concentrations assuming homogenous mixing of fluorescent : non-fluorescent LUV =1:7. Notably, this experiment showed a nearly constant $1/E_{FRET}$ throughout the F/T cycles, regardless of the initial phospholipid compositions, highlighting that the increase in $1/E_{FRET}$ observed in (A) reflects membrane mixing.

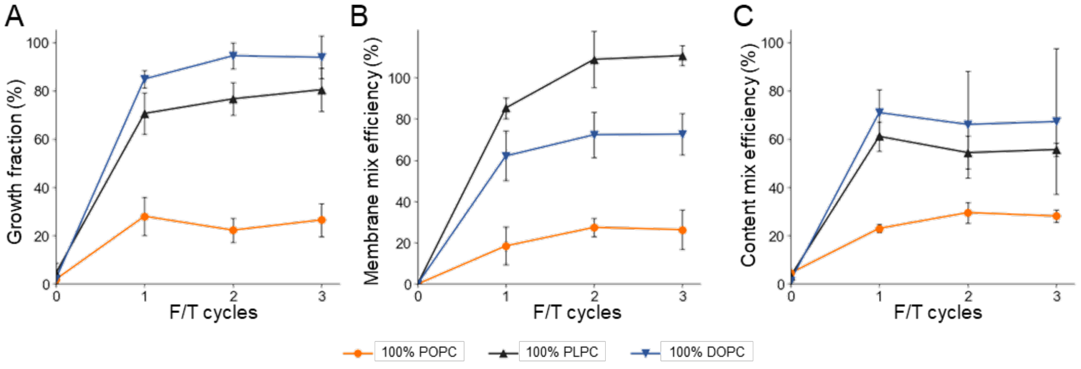


**Figure S7**: The result of (A) growth fraction, (B) membrane mix efficiency, and (C) content mix efficiency of 100% DOPC vesicles (blue). The data of 100% POPC (orange) and 100% PLPC (black) given in Figs 2–4, are shown for comparison. The vesicle composed of DOPC showed a higher value in every F/T cycle than POPC in all three features.

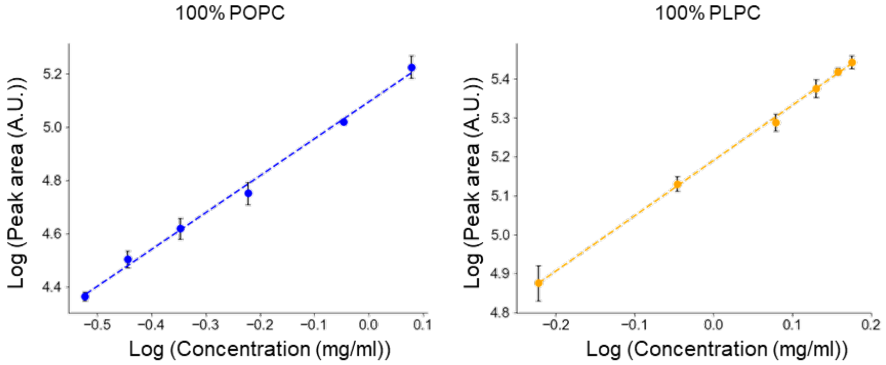


**Figure S8**: Standard curve used to quantify POPC and PLPC using HPLC/ELSD (Shimadzu Corp.). ELSD peak area attributed to POPC (retention time: ~5.3 min) and PLPC (retention time: ~4.3 min) versus their concentrations in the injected fluid of 5 $\mu$L were plotted. We confirmed a linear correlation between the logarithm of the concentration and that of the peak area for both POPC and PLPC.

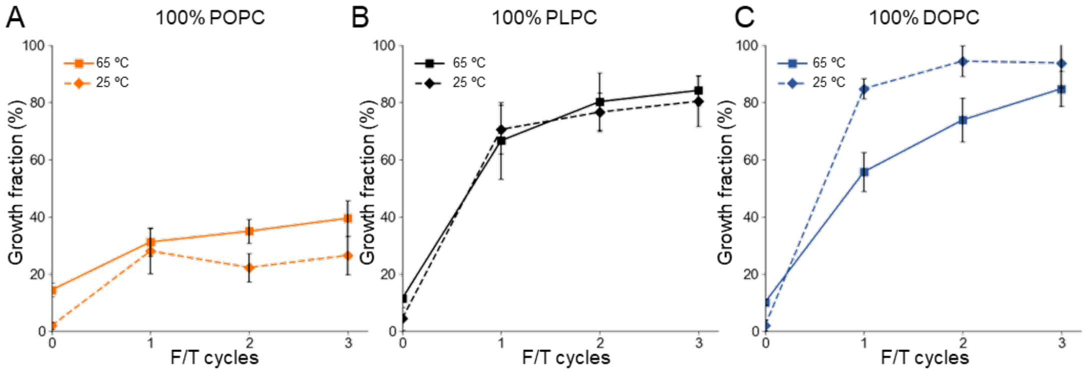


**Figure S9** Effect of thawing condition on the growth fraction of (A) 100% POPC, (B) 100% PLPC, and (C) 100% DOPC vesicles by F/T cycles. Here, vesicles were subjected to F/T cycles as described in the method section, with two different thawing conditions. The thawing was performed for around 10 min either at room temperature of approximately 25°C or 65°C in a dry bath incubator. All three vesicles showed no significant difference in growth fraction between thawing at 25°C and 65°C.
